# Activation of TAp73 and inhibition of thioredoxin reductase for improved cancer therapy in *TP*53 mutant pancreatic tumors

**DOI:** 10.1101/398750

**Authors:** Pilar Acedo, Aristi Fernandes, Joanna Zawacka-Pankau

**Affiliations:** Department of Microbiology, Tumor and Cell Biology, Karolinska Institutet, Biomedicum, Solnavägen 9, 141 8671 65, Stockholm, Sweden; Department of Medical Biochemistry and Biophysics, Karolinska Institutet, Biomedicum, Solnavägen 9, 171 65, Stockholm, Sweden

**Keywords:** protoporphyrin IX, benzoporphyrin derivative, TAp73, mutant p53, apoptosis, ROS, thioredoxin reductase

## Abstract

The p73 is a tumor suppressor that compensates for p53 loss and induces apoptosis in tumors in response to genotoxic stress or small-molecule treatments.

Pancreatic ductal adenocarcinoma (PDAC) has a late onset of the disease, responds poorly to the existing therapies and has very low overall survival rates.

Here, using drug-repurposing approach, we found that protoporphyrin IX (PpIX) and benzoporphyrin derivative monoacid ring A (BPD) activate p73 and induce apoptosis in pancreatic cancer cells. PpIX and BPD induce reactive oxygen species and inhibit thioredoxin reductase 1 (TrxR1). Thus, PpIX and BPD target cancer cells’ vulnerabilities namely activate TAp73 tumor suppressor and inhibit oncogenic TrxR1. Our findings, may contribute to faster repurposing of PpIX and BPD to treat pancreatic tumors.

**Lay Abstract:** Despite the efforts, pancreatic cancer remains among the most aggressive tumors. Late diagnoses often linked with the asymptomatic disease progression make it extremely difficult to cure. We have used drugs that are already in clinics and applied in photodynamic therapy of cancer and showed that the compounds induce death of cancer cells. The mechanism is via activation of p73 tumor suppressor and inhibition oncogenic thioredoxin reductase. Molecules that in parallel induce two pathways leading to cell death might be very promising candidates for improved cancer therapy in pancreatic cancer patients.

## Introduction

Pancreatic cancer is one of the deadliest diseases and the 5^th^ leading cause of cancer-related deaths worldwide[1]. In Europe, the death rates from pancreatic cancer continue to increase in both sexes, and the median survival rate for patients diagnosed with the metastatic disease is only 4.6 months[2]. At present, pancreatic cancer is the 4^th^ leading cause of death in Europe and is predicted to be the 2^nd^ cause of cancer-related deaths in 2020[3].

To date, it is still one of the most challenging solid tumors to treat. Most of the patients die within 1 year after diagnosis and only about 3-5% survive up to 5 years in the US. This is mainly due to the late onset of the disease, high incidence of metastasis and the lack of effective treatments[4].

Treatment of pancreatic cancer is difficult and depends on the stage of the disease at time of diagnosis. The response to the currently available therapies is disappointing and the poor outcome is likely stemming from a complex set of several factors described below.

In cases not suitable for resection with curative intent, palliative chemotherapy may be used to improve the quality of life. Gemcitabine (GEM) was approved by the Food and Drug administration in 1998 for the treatment of patients with advanced pancreatic cancer. GEM remains the main first-line therapy for pancreatic cancer patients regardless of primary and acquired resistance. The resistance is partially due to DNA-repair polymorphism[5], mutations in p53 pathway[6] or the existence of cancer stem cells[7]. Recently, combination treatment regimens with 5-fluorouracil, irinotecan and oxaliplatin (FOLFIRINOX) brought some promise to overcome treatment resistance in patients suffering from metastatic disease, however, pancreatic cancer remains a major unmet medical need[8].

Thus, there is an urgent need to develop more effective treatments for this type of cancer.

Pancreatic cancer is one of the cancer types in which mutant p53 impacts disease progression. *TP*53 is mutated in 75% cases of pancreatic carcinomas[9]. *TP*53 gene mutations co-exist with activating mutations in classical oncogenes including *K-Ras*, which accounts for 45% of all pancreatic cancer cases[10].

*TP*53 mutations are common across various tumor types and result with the loss of p53 tumor suppressor function and preferentially promote the so-called gain of new functions. As evidenced, mutant p53 is the main driver of disease progression in pancreatic cancers and selective therapies aiming at restoring p53 pathway to overcome p53 loss in cancer cells but not in normal cells, appear to be an attractive strategy for diseases such as chemotherapy-resistant pancreatic cancer. Despite the on going efforts, small molecules re-activating mutant p53 are still in clinical development[11].

It has been previously shown that photodynamic therapy of cancer (PDT), that uses photoactivatable compounds and light, is effective against numerous pancreatic cancer cell lines as monotherapy[12] or in combination with gemcitabine[13] or irinotecan[14].

The majority of the compounds that are clinically approved for PDT are porphyrins and their derivatives[15]. Protoporphryin IX (PpIX) is a metabolite of γ-aminolevulinic acid (ALA), a pro-drug applied in PDT of cancer[15]. Protoporphyrin IX, itself, without light excitation, was shown to induce wild-type p53-related cell death in several human cancer cell lines including human colon carcinoma[16]. Next, it has recently been demonstrated that PpIX stabilizes and activates TAp73 and induces TAp73-dependent apoptosis in cancer cells lacking *TP*53[17].

Thioredoxin reductase 1 (TrxR1) is a selenoprotein of the thioredoxin system and is central in maintaining redox homeostasis in cells[18]. TrxR has a protective role against oxidative stress and contributes to tumor progression once the disease has already developed. Thus, *TXNRD*1 is often overexpressed in cancers and is a promising target for improved cancer therapy[18].

Here, we found that PpIX and benzoporphyrin derivative (BPD; verteporfin®) induce TAp73 and its pro-apoptotic targets on mRNA and protein levels. Next, it was demonstrated that PpIX and BPD induce reactive oxygen species (ROS) and antioxidant response genes *HMOX*-1 and *NQO*1. PpIX and BPD inhibited TrxR1 *in vitro* and in pancreatic cancer cells harboring mutant p53. Taken together, our data adds up to the mechanism by which PpIX and now BPD induce apoptosis in cancer cells harboring *TP*53 gene mutations, which is via reactivation of TAp73 tumor suppressor and inhibition of TrxR1.

## Materials and methods

### Cell culture

Human pancreatic cancer cell lines, Paca3, MiaPaca2 and Panc1 were kindly provided by Dr. Rainer Heuchel from Karolinska Institutet, Sweden. Murine pancreatic cancer cells Panc02 were provided by Dr. Maximilian Schnurr from University of Munich, Germany and made available for this project by Dr. Yihai Cao from Karolinska Institutet, Sweden.

Normal pancreatic cells immortalized by HPV transformation (HPDE)[19] were provided by Dr. Rainer Heuchel from Karolinska Institutet, Sweden. Cells were maintained in DMEM medium supplemented with 10% fetal bovine serum and 1% penicillin/streptomycin solution (Sigma-Aldrich, Germany). The cells were cultured at 37°C in a humidified incubator with 5% CO_2_. The cell lines were routinely checked for mycoplasma during the whole duration of the project.

### Chemicals

Protoporphyrin IX and benzoporphyrin derivative were purchased from Sigma-Aldrich, dissolved in 100% DMSO (Sigma-Aldrich) to a final concentration of 2 mg/mL and stored in the dark in amber tubes at room temperature or at -20**°**C until use.

Cisplatin (CDDP) (Sigma-Aldrich) was dissolved in 0.9% NaCl to a concentration of 25□M and used at final concentration of 20 μM. Gemcitabine (2′,2′-difluoro-2′-deoxycytidine, GEM) (Abcam, UK) was prepared as 1 mM stock solution in 0.9% NaCl. Resveratrol (Enzo Biochem, USA) was reconstituted to 40 mM in 100% DMSO, aliquoted and stored at -20°C.

MTT reagent (Sigma-Aldrich, Germany) was reconstituted in PBS (prepared from 1x tablets Sigma-Aldrich, Germany) to 5 mg/ml, filter-sterilized and stored at +4°C for a week.

### Verteporfin uptake and cellular localization

Cells were incubated with 2.5 μg/ml of PpIX or BPD in cell media for 3h, washed three times with PBS and co-stained with ER-Tracker™ Green for live-cell imaging (Molecular probes) according to manufacturer protocol. DAPI (Molecular Probes) was used for nuclear counterstaining. Organelle marker was excited using a 488 nm laser and BPD was excited with 420nm (emission at 690 nm). Images were taken using an Olympus IX-71 microscope and DeltaVision SoftWoRx.

### Cell viability

Cell viability was assessed by MTT assay as provided by the manufacturer (Sigma-Aldrich, Germany) and as described previously[16]. Briefly, 3000 cells were seeded in 96-well plate and allowed to adhere overnight. The cells were treated with increasing concentrations of compounds for 72 h and the viability was measured using MTT reagent.

### Real-time PCR

Quantitative PCR was performed as described previously[20]. Cells were seeded in 6-well plates, allowed to adhere for 24 h and treated with 2.5 μg/ml PpIX or BPD for 8h. mRNA after isolation was reverse transcribed to cDNA according to the manufacturer’s instructions (Bio-Rad, Sweden). For qPCR reaction 150□nM; 10□ng cDNA; 7.5□μl 2 × master mix (Bio-Rad) and water to a total of 15□μl were used. Primers used: *GAPDH* forward: TCATTTCCTGGTATGACAACG and reverse: ATGTGGGCCATGAGGT *, NOXA*(PMAIP1) forward 5′-AAGTGCAAGTAGCTGGAAG-3′, reverse: 5′-TGTCTCCAAATCTCCTGAGT-3′, *PUMA* forward: 5′-CTCAACGCACAGTACGAG-3′ and reverse: 5′-GTCCCATGAGATTGTACAG-3′, *HMOX-1* forward: 5′-TTCACCTTCCCCAACATTGC-3′ and reverse: 5′-TATCACCCTCTGCCTGACTG-3′, *Bid* forward: 5′-GTGAGGTCAACAACGGTTCC-3′ and reverse: 5′-TGCCTCTATTCTTCCCAAGC-3′, *Bim* forward: 5′-TGGCAAAGCAACCTTCTGATG-3′ and reverse: 5′-GCAGGCTGCAATTGTCTACCT-3′, *Bax* forward: 5’ – GCTGTTGGGCTGGATCCAAG – 3’ and reverse: 5’ – TCAGCCCATCTTCTTCCAGA – 3’

### Thioredoxin reductase activity assay

For the enzymatic activity of purified TrxR1 we followed the protocol previously described[21, 22]. Briefly, 10 nM of purified TrxR1[23] was pre-incubated with BPD at the indicated concentrations (0–20 μM) in 50 μl PE-buffer supplemented with 150 μM NADPH for 30 min at room temperature. Next, 2.5 mM DTNB and 150 μM NADPH were added to the samples just before measurement. DTNB reduction was performed for 6 min by detection of TNB anion formation detected as a change in the absorbance at 412 nm.

For cellular TrxR activity Panc1 and MiaPaCa2 cells were treated with 2.5 μg/mL PpIX or BPD for 6 h after which they were harvested and re-suspended in TE buffer containing a protease inhibitor cocktail (cOmplete, mini protease cocktail tablets, Roche, Sweden). Cells were sonicated and the total protein concentration of the supernatant was determined using a Bradford reagent kit (Bio-Rad Laboratories, Sweden). Cellular TrxR1 activity was determined using the previously described end-point Trx-dependent insulin reduction assay[22]. Briefly, total cellular protein (20 μg) was incubated with 15 μM recombinant human wild-type Trx in the presence of 297 μM insulin, 1.3 mM NADPH, 85 mM Hepes buffer, pH 7.6, and 13 mM EDTA for 30 min at 37°C, in a total volume of 50 μl. The reaction was stopped by the addition of 200 μl of 7.2 M guanidine–HCl in 0.2 M Tris–HCl, pH 8.0, containing 1 mM DTNB. The activity was then determined by measuring absorbance at 412 nm using a VersaMax microplate reader (Molecular Devices, USA) with a background absorbance reference for each sample containing all components except purified TrxR, incubated and treated in the same manner.

### Cell death detection

Propidum iodide and FITC-Annexin V (from BD Biosciences, USA) staining was performed according to the manufacturer’s protocols and FACS analysis was performed using the CELLQuest software (CELLQuest, Franklin Lakes, NJ, USA) as described previously[24].

### ROS measurement

Reactive oxygen species were detected using 2’,7’-dichlorofluorescein (H_2_DCF) (Sigma-Aldrich, Germany) and method described previously[24].

Hydroethidine staining was performed according to the manufacturer protocol (Life Technologies, Sweden). Data analysis was performed with the CELLQuest software (CELLQuest, Franklin Lakes, NJ, USA).

### Western blotting

Western blotting was performed according to the standard protocol. Briefly, 100□μg of total cell lysate was subjected to electrophoresis, after transfer the following antibodies were used to detect proteins: anti-TAp73 (Bethyl Laboratories, TX, USA), anti-BAX (Santa Cruz Biotechnology, Germany) anti-PUMA (Merck, MA, USA), anti-HO-1 (Santa Cruz Biotechnology), anti-NRF2 (Santa Cruz Biotechnology, Germany), anti-Bid (Santa Cruz Biotechnology), and anti-actin (Sigma-Aldrich, Germany), anti-PARP (Santa Cruz Biotechnology, Germany), anti-actin (Sigma-Aldrich, Germany).

## Results

### Benzoporphyrin derivative localize into the cytoplasm of Panc1 cancer cells

Pancreatic ductal adenocarcinomas (PDAC) are cancers of poor clinical outcome due to chemoresistance and disease relapse and mutant p53 drives disease aggressiveness. Whether the re-activation of TAp73 can compensate for the p53 loss in cancers with *TP*53 gain of function mutations has not been unequivocally demonstrated yet. For this reason, we tested the outcome of TAp73 activation in PDAC cancer cell lines harboring hot-spot *TP*53R273H mutation.

First, we assessed the subcellular localization of BPD in Panc1 cells (Fig. 1a). Fluorescence microscopy indicated that BPD (red fluorescence emission) is uptaken to high degree by cancer cells. Next, we found that verteporfin localized in large in cytoplasm and partially in endoplasmic reticulum.

**Figure 1.**
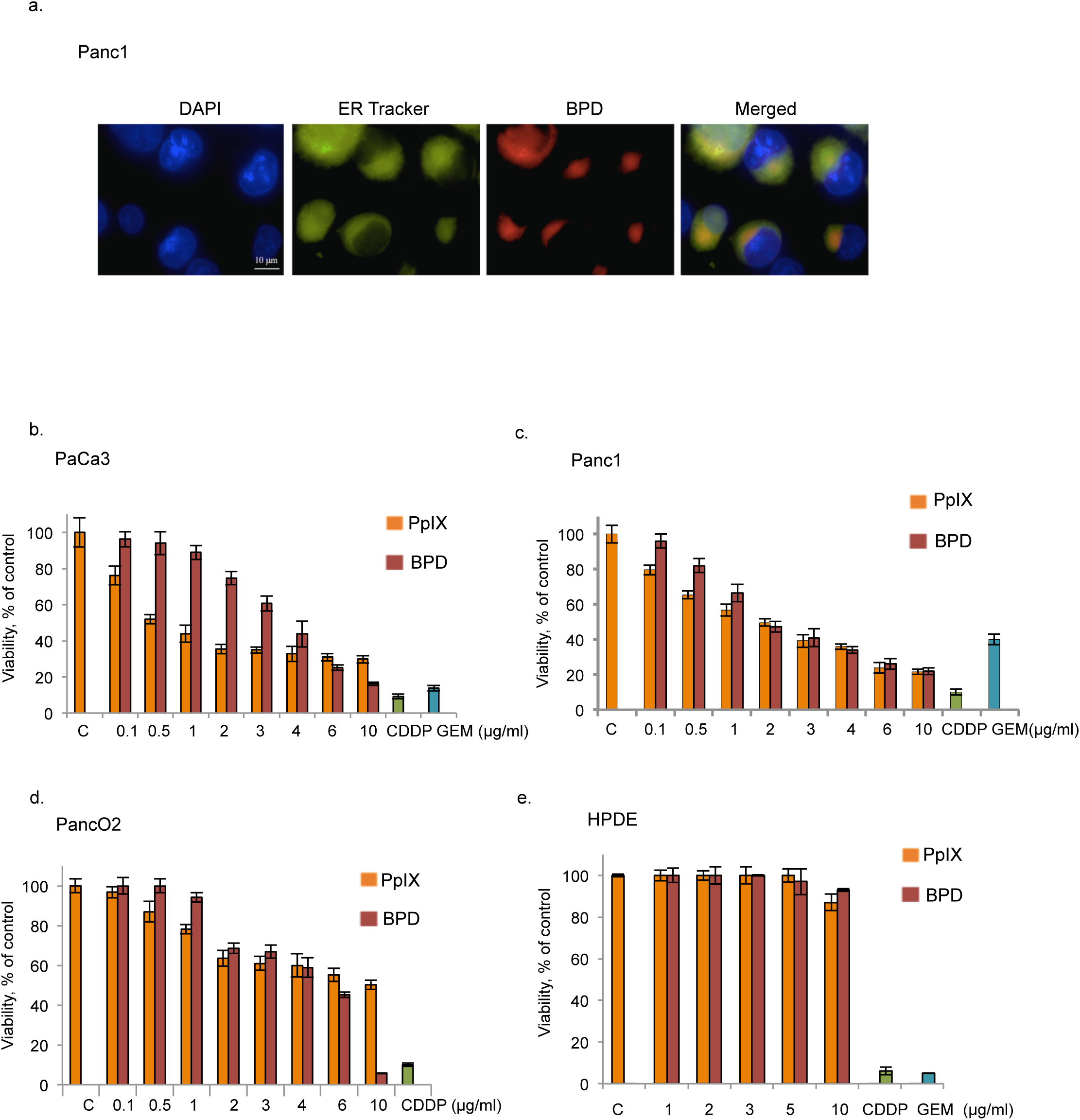
BPD accumulates and inhibits proliferation of pancreatic cancer cells. (a) BPD accumulates largely in the cytoplasm and partially in endoplasmic reticulum in Panc1 cancer cells. Nuclei were stained with DAPI and endoplasmic reticulum with ER-tracker Green, respectively. The image is a representative data of three independent experiments. (b) PpIX (orange bars) and BPD (red bars) inhibit proliferation of PaCa3, Panc1 (c) and Panc02 (d) cancer cells as assessed by MTT assay after 72 hours treatment. 25 μM cisplatin (CDDP, green bars) and 5 μM gemcitabine (GEM, blue bars) were used for comparison. (e) PpIX (orange bars) and BPD (red bars) do not inhibit proliferation of HPDE cells as assessed by MTT assay.

### PpIX and BPD inhibit proliferation of PDAC cells but not of non-transformed human ductal epithelial cells

To assess the impact of PpIX and BPD on the proliferation of PDAC cells we performed MTT proliferation assay. Standard treatments for advanced pancreatic cancer disease include monotherapy with gemcitabine (GEM) and we used it for comparative purpose.

MTT assays showed that PpIX and BPD inhibit cancer cells’ proliferation (Fig. 1b-e). In agreement with previous data, cancer cells expressing wild-type (wt) p53 (PaCa3) were sensitive to 72h-treatment with PpIX[16] (Fig. 1b) and showed enhanced growth inhibition upon BPD exposure. Interestingly, mutant p53-harboring cells (Panc1) were more sensitive to PpIX than PaCa3 cells and slightly less sensitive to BPD (Fig. 1c).

The calculated IC_50_ for PpIX was 4.4 µM for PaCa3 (wtp53) cells and 3.5 µM for Panc1 (mtp53R273H) and 2.2 and 2.8 µM for BPD, respectively.

Of note, murine pancreatic cancer cells Panc02 and human MiaPaCa2, were less sensitive to PpIX and BPD at the range of the concentrations tested (Fig. 1d and data not shown) and normal human epithelial ductal cells were insensitive to the concentrations tested (Fig. 1e). On other hand, all cell lines were highly susceptible to low dose (25 µM) cisplatin and 5 µM gemcitabine.

Consistent with previous data wtp53 cancer cells were sensitive to PpIX[16]. Interestingly, mutant p53 Panc1 cell line was the most sensitive among the cells tested. Based on the above and previous studies, we selected Panc1 cell line and 2.5µg/ml compounds’ concentration for further studies on cell death induction upon reactivation of TAp73.

### PpIX and BPD induce apoptosis in PDAC cells

To investigate if the growth inhibition observed in MTT assay was a consequence of the induction of apoptosis, we treated PaCa3 and Panc1 cells for 48 h with 2.5 µg/ml PpIX or BPD and performed flow cytometry analysis using propidium iodide (PI) and Annexin V stainings. We found that at the time-point tested, Panc1 undergo high late apoptosis after PpIX treatment as demonstrated by above 40% increase of PI positive cells relative to untreated control samples. At this time-point, Annexin V positivity, indicating early apoptosis, increased by 12% or 9% after PpIX or BPD treatments, respectively (Fig. 2c,d). In PaCa3 cells, harboring wtp53, we observed 10% and 17% increase in PI positive cells after PpIX or BPD treatment, respectively. The increase in Annexin V-positive cells was 5% for PpIX and 11% for BPD. Thus, both compounds induce early and late apoptosis in cancer cell lines tested. Cisplatin treatment was used as a positive control in these assays.

**Figure 2.**
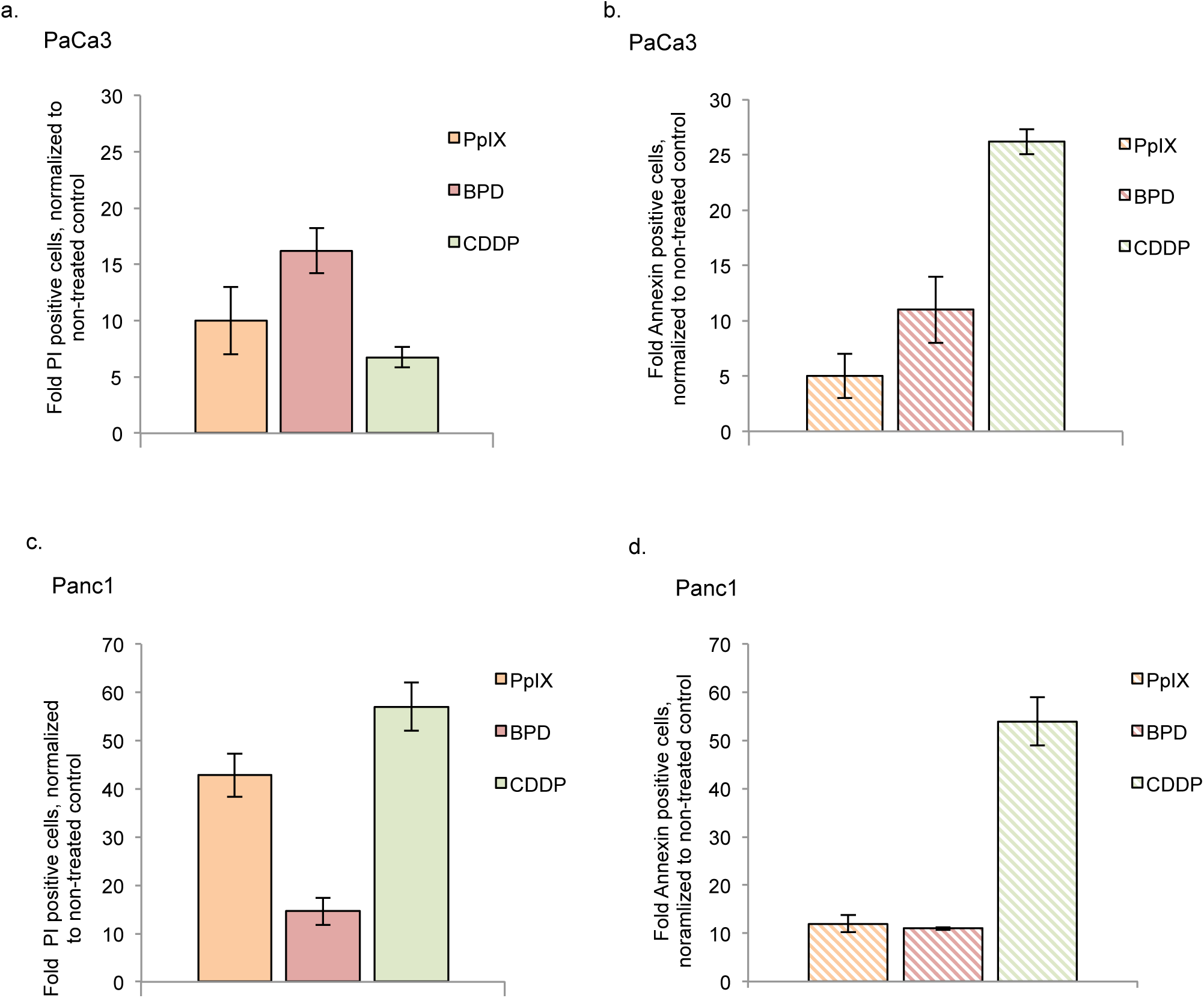
PpIX and BPD activate early and late apoptosis in PDAC. (a) Treatment with 2.5 μg/ml PpIX (orange bars) and BPD (red bars) induced late (PI staining) and early apoptosis (FITC-Annexin V) (b) in wtp53 PaCa3 cells after 48 hrs (n=3). (c) Treatment with 2.5 μg/ml PpIX and BPD for 48 hrs induced late (PI staining) and early apoptosis (FITC-Annexin V) (d) in mtp53 Panc1 cells (n=3). 20 μM CDDP was used as a positive control for apoptosis detection.

### PpIX and BPD induce TAp73 and its pro-apoptotic targets

It has been demonstrated that PpIX reactivates TAp73 tumor suppressor and induces TAp73-dependent apoptosis in *TP*53-null cancer cells[17]. To assess if the growth inhibition observed in PDAC cell lines by PpIX and BPD is a result of apoptosis, we evaluated the expression of TAp73 and its pro-apoptotic targets in two pancreatic cancer cells lines. Western blotting results showed that in dark conditions, both PpIX and BPD induce TAp73 levels, which correlates with the accumulation of its proapoptotic Bcl-2 protein family members (PUMA, Bax, and Bid) (Fig. 3a,b,c) and cleaved PARP in a time dependent manner. In Panc1 cells PpIX and BPD upregulate the expression of bax, bid, puma and noxa (Fig. 3d). This indicates that cell apoptosis is due to activation of TAp73 (Fig. 3a, c). Interestingly, we detected activation of antioxidant response proteins such as HO-1, and NF-E2-related transcription factor (Nrf2) (Fig. 3a,b,c,d). PpIX and BPD did not induce PARP cleavage or activation of TAp73 and its proapoptotic targets in HPDE cells at the effective concentrations (Fig. 3e).

**Figure 3.**
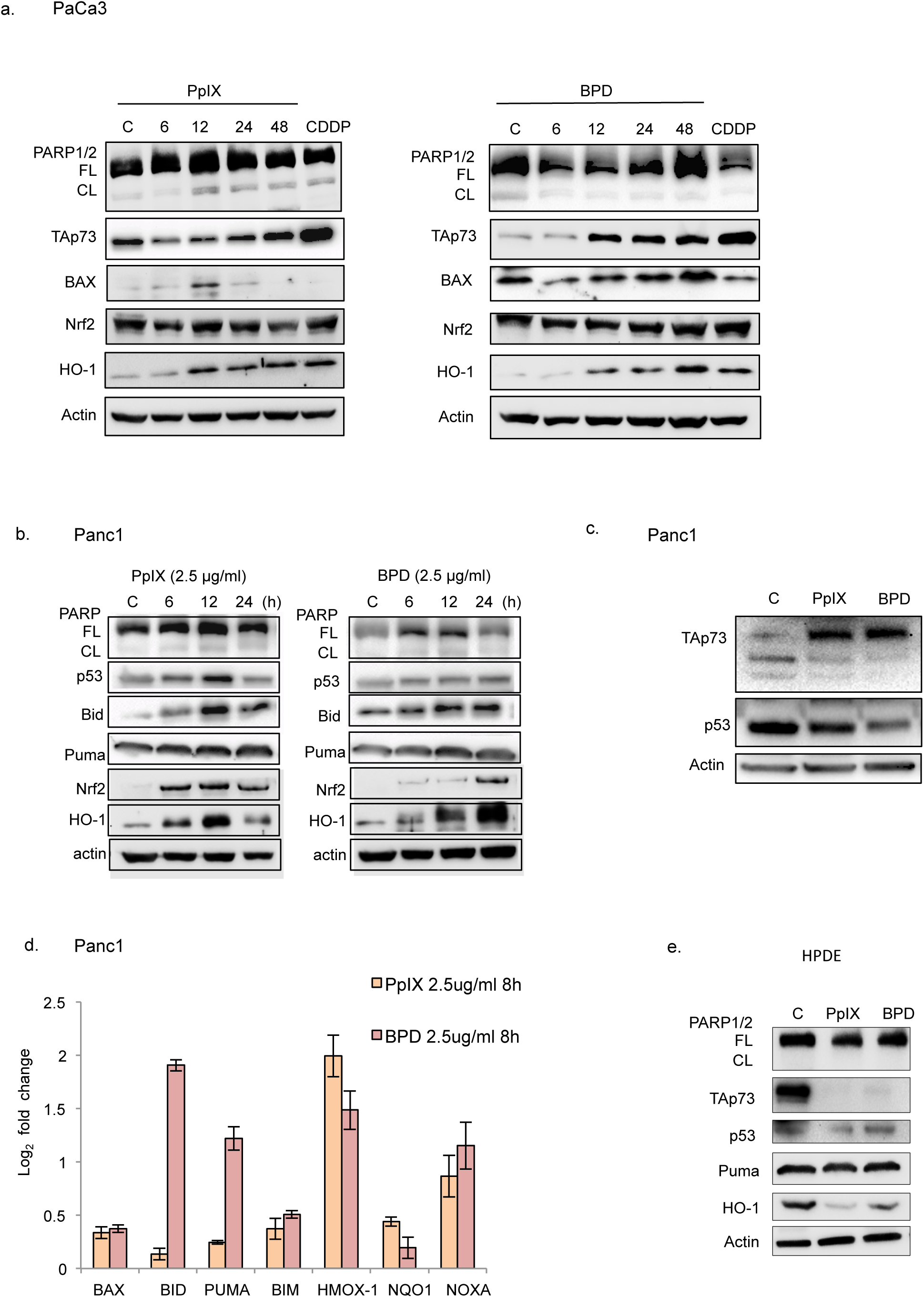
Reactivation of TAp73 by PpIX and BPD induces apoptosis in pancreatic cancer cells. (a) PpIX and to lower extend BPD (2.5 μg/ml) induce PARP cleavage in PaCa3 and Panc1 cells (b, c). The effect is time dependent. The PARP cleavage correlates with stabilization of TAp73 protein levels and its pro-apoptotic target Bax. Both compounds upregulated heme oxygenase (HO-1) and slightly its transcriptional regulator Nrf2 protein. 20 μM CDDP was used as a positive control. FL – full length; CL – cleaved. (d) Treatment with PpIX (orange bars) and BPD (red bars) increased mRNA levels of apoptotic proteins and antioxidant response proteins in Panc1 cells. (e) PpIX and BPD treatment did not induce TAp73 or apoptotic proteins in non-transformed pancreatic cells.

### PpIX and BPD induce reactive oxygen species in PDAC

We observed accumulation of antioxidant response proteins after treatment with PpIX and BPD in PDAC cells; we thus sought to investigate if PpIX and BPD trigger accumulation of reactive oxygen species (ROS). We assessed the levels of ROS using DCFDA and HE probes. Fig. 4a,b,c shows that PpIX and BPD induce ROS in PaCa3 and Panc1 cancer cells. Pre-treatment with the ROS scavenger resveratrol inhibited the generation of ROS by PpIX and BPD in Panc1 cells (Suppl. Fig. 1a). Importantly, the compounds did not induce ROS in HPDE cells (Suppl. Fig. 1b).

**Figure 4.**
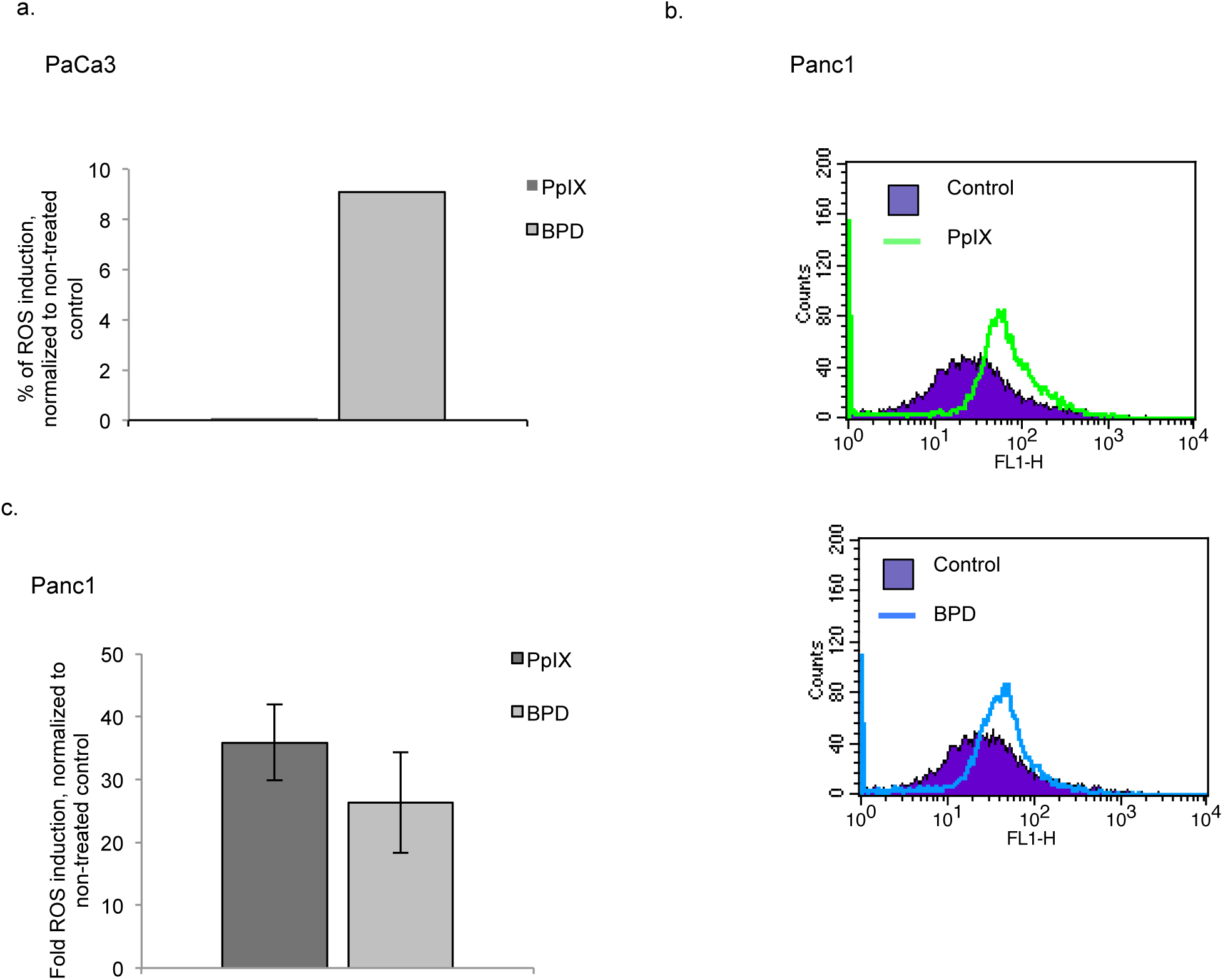
PpIX and BPD induce ROS in pancreatic cancer cells. (a) BPD significantly increased ROS levels in PaCa3 cells as estimated by DCFDA measurement 16 h post-treatment (n=3). (c) Both, PpIX and BPD, induced ROS in Panc1 cells as assessed by DCFDA staining (n=3).

### PpIX and BPD are inhibitors of thioredoxin reductase

Protoporphyrin IX was first identified as an inhibitor of thioredoxin reductase in an *in vitro* screen[25]. Since we observed significant induction of ROS in cells treated with PpIX or BPD and inhibition of TrxR is known to cause induction of ROS, we reasoned that apart from active apoptosis, inhibition of the enzyme might contribute to elevation of ROS in cancer cells.

As demonstrated in Fig. 5a, recombinant TrxR was inhibited by both, PpIX and BPD. To assess if PpIX and BPD inhibit TrxR activity in cells, we treated Panc1 with PpIX or BPD for 6 h and assayed the activity using end-point Trx-dependent insulin reduction assay. PpIX and BPD treatment inhibited TrxR1 in Panc1 cells (Fig. 5b) and in other mutant p53 harboring cell line, MiaPaCa2. Taken together, these data demonstrate that BPD, similarly to PpIX, is a potent inhibitor of TrxR in pancreatic cancer cells.

**Figure 5.**
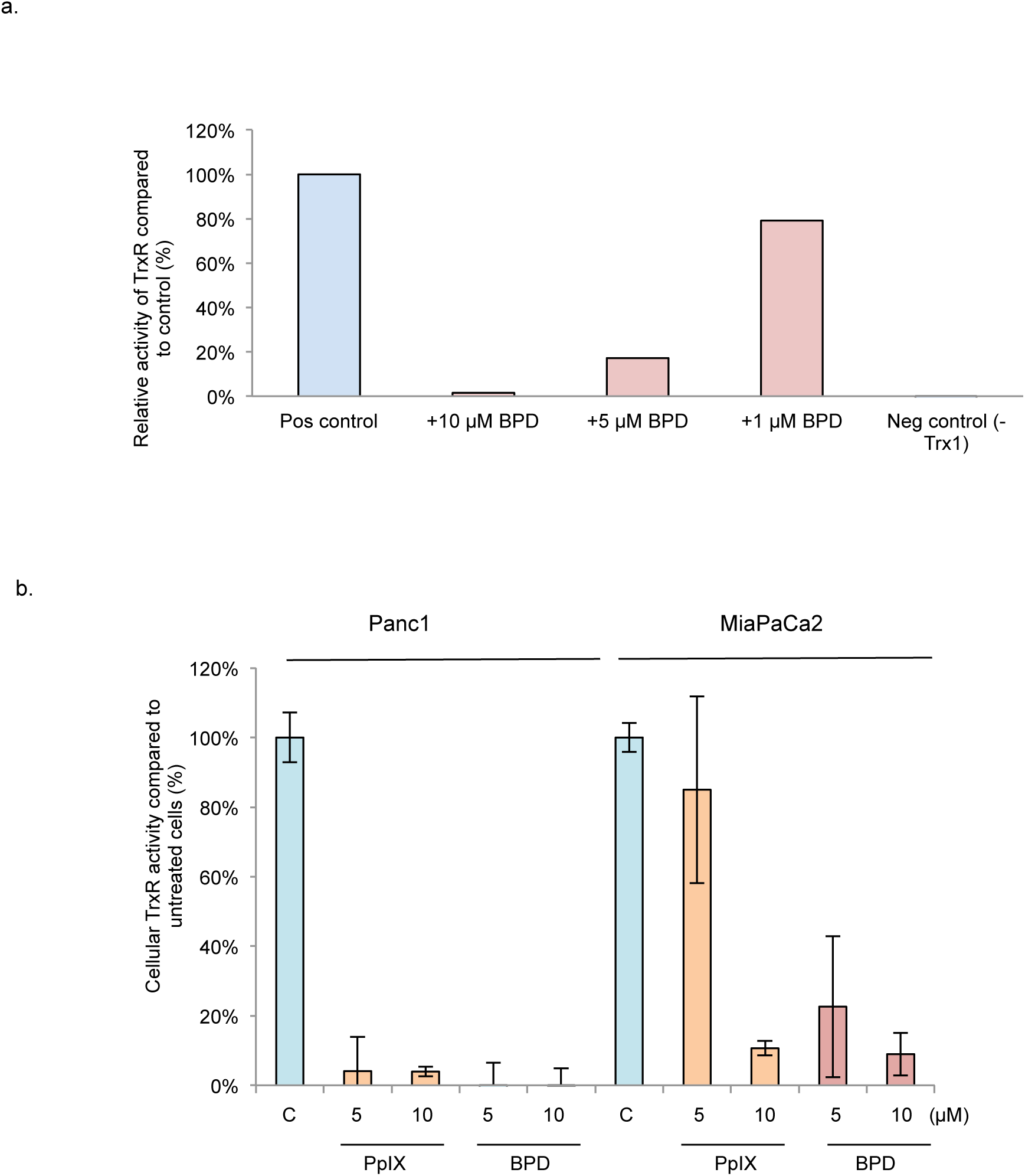
PpIX and BPD inhibit thioredoxin reductase activity. (a) BPD (red bars) ablates activity of recombinant TrxR. Representative data of three independent experiments is presented. (b) Both, PpIX (orange bars) and BPD (red bars) inhibit TrxR in Panc1 and MiaPaCa2 pancreatic cancer cells (n=2).

## Discussion

Several protein kinases inhibitors were reported to have off-target effects. In example, BRAF inhibitor, vemurafenib, used for treating metastatic melanoma[26] with good clinical outcome in some cases[27], was manifested to induce unusual photosensitivity in patients[28]. A recent study by Klaeger *et al*. [29] demonstrated that several clinically approved kinase inhibitors, including vemurafenib, in addition to their designed targets effectively inhibit ferrochelatase, the last enzyme in heme biosynthesis. The authors speculated that accumulation of PpIX may lead to photosensitivity of the patients upon treatment with kinase inhibitors.

Next, PpIX is a metabolite of γ-aminolevulinic acid (ALA), heme precursor, a drug applied in photodynamic therapy of cancer [15].

We have shown in a drug-repurposing approach that PpIX re-activates wild-type p53 and TAp73 and induces apoptosis in cancer cells. The mechanism is *via* inhibition of p53/MDM2 and TAp73/MDM2(X) interactions[16, 17]. Next, PpIX was identified in the *in vitro* screening as a potent, competitive inhibitor of thioredoxin reductase (TrxR)[25]. Thus, here we speculated that PpIX, exogenously delivered to pancreatic cancer cells with *TP*53 mutations, will induce apoptosis *via* activation of p53 protein family member and may simultaneously inhibit TrxR.

The detailed analysis revealed that PpIX and its clinically applied analog, BPD, known under the commercial name of verteporfin®, inhibit proliferation of pancreatic cancer cells without affecting non-transformed cells. The mechanism of growth inhibition was by induction of apoptosis. Next, both compounds triggered antioxidant response and induced accumulation of ROS only in cancer cells but not in normal HPDE cells. The above, prompted us to investigate if robust induction of ROS is a consequence of inhibition of TrxR. According to our findings, BPD inhibited enzymatic activity of recombinant thioredoxin reductase. Both, PpIX and BPD inhibited this enzyme in Panc1 and MiaPaCa2 cancer cells as demonstrated by Trx-dependent end point insulin reduction assay.

Inhibitors of TrxR are currently under pre-clinical development and are promising candidates for cancer treatment[30]. Several small molecules activating wild-type or mutant p53 were also shown to inhibit TrxR[21,31,32]. The small molecule APR-246, APR-246, a mutant p53-reactivating compound in Phase II clinical development, is a pro-drug, converted in cells to methylene quinuclidinone (MQ) [33]. MQ is a Michael acceptor and was demonstrated to target cysteines in p53 core domain [34] and selenocysteine (Sec) residues in the C-terminal motif of TrxR1[31]. Our findings clearly demonstrate that PpIX and BPD inhibit TrxR. The mechanism by which this occurs remains to be elucidated. Structural studies on heme-binding protein, heme oxygenase 2 (HO-2), revealed that in the oxidized state of cysteine residues in apoprotein, heme binds 2.5 fold more tightly than in the reduced state[35]. This, together with the finding that the Sec-to-Cys mutant of TrxR1 was resistant to inhibition by PpIX[25], strongly imply that Sec residue in TrxR might be the binding site of PpIX and BPD (verteporfin®).

In conclusion, we showed that PpIX and now BPD activate TAp73 in pancreatic cancer cells, which results in apoptotic cell death. In parallel the compounds induce ROS, which contributes to cell killing. Induction of high ROS is most probably due to inhibition of TrxR by PpIX and BPD.

### Future perspective

Our findings might have important clinical implication and support fast repurposing of porphyrins in treating pancreatic cancer patients harboring *TP*53 mutations. Firstly, the reactivation of TAp73 and inhibition of TrxR might help to predict the response of patients to PpIX or BPD. Next, dual targeting of TAp73 and TrxR might bring additional advantage in overcoming the development of treatment-resistant disease. In particular, we foresee that parallel reactivation of TAp73 and inhibition of TrxR will significantly increase the stress burden in cancer cells when compared to separate approaches, which will contribute to selective tumor killing without affecting normal cells.

### Executive summary

- PpIX and BPD induce apoptosis in PDAC cells by activating TAp73
- PpIX and BPD induce reactive oxygen species in PDAC by inhibiting thioredoxin reductase

---

**Supplementary Figure 1. PpIX and BPD induce ROS generation.**

(a) PpIX and BPD generated ROS in Panc1 cells as assessed by HE staining. Pre-treatment with 4 μM resveratrol reverted the effect of these compounds. Representative histograms of three independent experiments are shown.

(b) Neither, PpIX nor BPD induced ROS in non-transformed HPDE cells (DCFDA staining). H_2_O_2_ treatment was used as positive control of ROS induction. Representative histograms are shown of three independent experiments.

## Acknowledgements

Special thanks are addressed to Dr. Alicja Sznarkowska from University of Gdansk for helpful discussions. We would like to thank all our colleagues for sharing their reagents.

Author Contributions
P.A. performed experiments, drafted figures, read the final version of the manuscript A.F. participated in designing the experimental work for thioredoxin reductase measurement; read the final version of the manuscript
JZP designed the study, obtained the funding, selected the methods, drafted the manuscript, prepared final figures, prepared the final version of the manuscript, was responsible for correspondence with the Journal.

## Notes

1 **Financial disclosure:** The work has been supported by Karolinska Institutet/MDAnderson Cancer Center Sister Institution Funds and The Strategic Research Programe in Cancer, Stockholm Läns Landsting and Åke Wibergs stiftelse.

